# A recombinant baculovirus TnGV in *Trichoplusia ni* larvae using the PIG bombardment method and the CRISPR/Cas9 system

**DOI:** 10.1101/2021.08.01.454629

**Authors:** Oscar J. Ortiz-Arrazola, Ma. Cristina Del Rincón-Castro

## Abstract

Baculoviruses have been used for the expression of heterologous proteins of biotechnological interest. However, most of these proteins are obtained by homologous co-transfection recombination in cell lines, limiting their use. Recently, the CRISPR/Cas9 system has excelled in its high efficiency in editing specific sequences without the need for insect cell lines. In this work, the CRISPR/Cas9 system was used to edit the genome of *Trichopusia ni* granulovirus (TnGV) and transformation of insects by the PIG bombardment method. A homologous repair vector (pTnGV101) was designed with regions *orf5* and *orf7*, as well as sgRNA flanking TnGV P10 of this virus. The bombardment transformation was carried out at 175 psi with 40% of infected *T. ni* larvae, of which 38% expressed the reporter protein EGFP. These results demonstrate that the CRISPR/Cas9 system and PIG bombardment can be used for genetic modification of baculovirus *in vivo*.

## Introduction

Baculoviruses are an important group of entomopathogens that are used to control a wide variety of pest insects (Szewczyk et al., 2006). The Baculoviridae family includes 4 genera: *Alphabaculovirus* (Nucleopolyhedrovirus or NPV), *Betabaculovirus* (Granulovirus or GV), *Gamabaculovirus* (NPV) and *Deltabaculovirus* (NPV). The baculovirus genome is a double-stranded supercoiled DNA of 82-180 kb (Jehle et al., 2006). Baculoviruses have been widely used for the expression of heterologous proteins (Airenne et al., 2013; Kost et al., 2005). Recombinant strains have been developed, inserting genes into the genome of some baculoviruses with the aim of increasing their virulence. Examples of this strategy are gene insertions coding for scorpion neurotoxins (Carbonell et al., 1988; Stewart et al., 1991), mite toxins (Tomalski, & Miller, 1991), the diuretic hormone of insects (Maeda, 1989), the sterease of the juvenile hormone (Hammock, et al., 1990), the venom of spiders, genes of anemones (Prikhod’ko et al., 1996) and for the *cry* genes of *Bacillus thuringiensis* (Martens et al., 1990; Merryweather et al., 1990).

Recombinant baculoviruses have only been developed in *Autographa californica* multiple nucleopolyhedrovirus (AcMNPV), and in *Bombix mori* nucleopolyhedrovirus (BmSNPV) (Kost et al., 2005; Martínez-Solís & Herrero, 2019; Motohashi et al. 2005). Most genetically modified baculoviruses have been obtained by homologous recombination by co-transfection of viral DNA and a transfer vector in cell lines; or the bacmids system for genetic exchange. Both transfer systems are limited by the few insect cell lines available (Hajós et al., 1998; Luckow et al., 1993). On the other hand, the CRISPR/Cas9 system has been successfully used for the deletion and insertion of genes in cell lines and higher organisms (Esvelt & Church, 2013; Lin et al., 2016). This method is based on the combination of a DNA endonuclease and a single RNA guide molecule (sgRNA) that directs the nuclease to a complementary target in DNA where it induces double-stranded breaks. In most cases, these lesions are repaired by non-homologous end-union (NHEJ) or homology-directed repair (HDR) that uses a homologous DNA template as a guide (Ran et al., 2013). The bombardment particle release system (originally used to transform plant cells) (Klein et al., 1988) uses tungsten or gold microparticles that are coated with DNA. These microparticles are inserted into the cell core at high speed. Are currently two types of particle bombardment systems in genetic transformation: the so-called Bio Rad PDS 1000 / HE (drying system) and the particle flow gun (PIG) (wet system) (Finer et al., 1992). The PIG system uses a dry layer of DNA-coated particles on a membrane, which is subjected to a sudden helium trigger. The membrane impacts a metal grid releasing the particles at a high velocity which penetrate the tissues of the white organism (Gunadi et al., 2019; Thomas et al., 2001).

The principal objective of this work was to use the CRISPR/Cas9 system to edit the *Trichoplusia ni* granulovirus (TnGV) genome in combination with the PIG bombardment technique to obtain a recombinant virus. A repair vector was developed which, when evaluated, showed the potential of baculoviruses to produce recombinant proteins when they do not have insect cell lines for genetic modification.

## Materials and methods

### Insects

A colony of *Trichoplusia ni* was established under insectary conditions at 28° C ± 2° C, with photoperiod of 14 hours day and 10 hours night and 60-70% RH. The larvae were fed in modified artificial semi-diet (Guy et al., 1985).

### Strains and vectors

The baculovirus strain used in this study was the granulovirus (TnGV) isolated from *Trichoplusia ni* and the transfer vector pAcUW31 (CLONTECH Laboratories Inc. Palo Alto, CA) was used as a template repair plasmid. The plasmid pU6-BBSI (donated by Izuho Hatada, Addgene plasmid #128433) was used for sgRNA cloning. Plasmid vector pHSP70-Cas9 (donated by L. B. Voshall, Addgene #45945) (Gratz et al., 2013) containing protein Cas9 under temperature induction promoter HSP70 of *D. melanogaster* was used as endonuclease. All plasmids were amplified in the *E. coli* TOP10F strain (Invitrogen Life Technologies, Carlsbad, CA).

### Construction of the repair vector

For the construction of the repair vector, it was used as template pAcUW31, to which the homologous regions ORF*lef2*/*orf603* and *orf1629* were changed to regions *orf5* and *orf7* of the wild TnGV baculovirus. These regions were amplified with oligonucleotides D-ORF5 5′CCGcggagCAAGGGTAGCCTGTCTATTGGAC 3′ and R-ORF5 5′ ATAAGAATGATCATCGGATTGGTGAAATTTTTCG 3′ for a fragment of 855 bp; and with D-ORF7 5′ATAAGAATCAGATCAGATG CCGctcgagTTCTTCATCTTCAATATTATGTC3 for a fragment of 3′ and 803-ORF7 5′. Once the repair recombination plasmid was constructed ( pTnGV101), the *egfp* reporter gene was inserted under promoter P10.

### Design and cloning of RNA guidance

The sgRNA were designed on the CHOP-CHOP platform (Labun et al., 2019) and synthesized by the commercial house T4 OLIGO. The sgRNA design was based on TnGV region P10 protein, selecting the sgRNA with the highest score within the white region (Table 1). For the assembly of sgRNA, 1 μl of each oligonucleotide was used incubating at 95° C for 5 minutes and then cloned in the pU6-BBSI vector. The resulting modified vector was named pU6-BBSI-grna. To verify the cloning of the sgRNA in the pU6-BBSI vector, the cloning site was amplified between T7 and T3 (700-850pb) of the pU6-BBSI-grna vector, with the initiators T7 5′TAATACGACTACTATAGGG and T3 5′GCAATTAACCCTACTAAAGG. The amplified fragments were sequenced at Macrogen Inc.

**Table 1.**
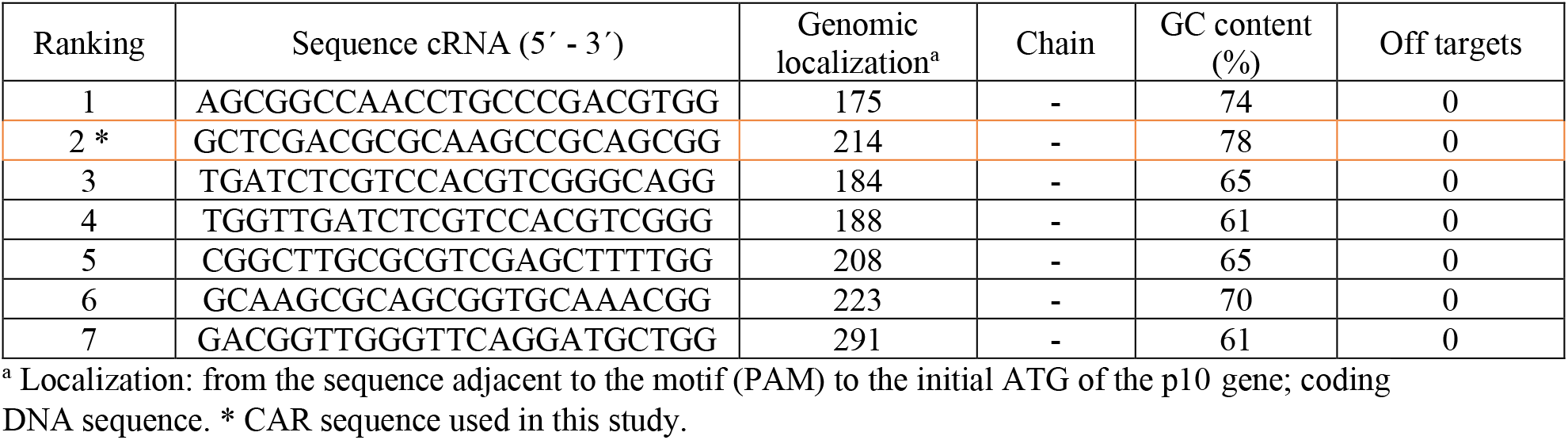
sgRNA designed on the CHOP-CHOP platform.

### Viral DNA extraction

TnGV purified virions were resuspended in 400 μl proteinase K buffer (Tris-HCl 0.01 M, EDTA 0,005 M, SDS 0.5%) and incubated at 60°C for 15 minutes. Subsequently, 100 μl of proteinase K (200 μg/ml) was added and incubated at 37°C for 2 h. DNA extraction was done with phenol:chloroform:isoamyl alcohol (25: 24:1) followed by chloroform:isoamylic alcohol (24:1). The mixture was fractionated by centrifuging at 10.000 g per 5 min. The samples were dialyzed all night using Tris/EDTA buffer. Finally, DNA was stored at −20°C until use in pig co-transfection.

### Preparation of gold particles for bombardment with the system PIG

Gold particles of ~ 1 μm were used (BioRad, Hercules, USA). USA). The particles were prepared at a concentration of 60 mg/ml as described by Cabrera-Ponce et al., (1997). The shooting pressures used were 160 and 175 psi. For each shoot pressure, three treatments were used with different DNA concentrations of vectors pTnGV101, pU6-BBSI-RNA, pHSP70/Cas9 and of the wild strain TnGV (6 treatments in total) (Table 2) resuspended in TE buffer (Tris-HCl 10 mm, 1 mm EDTA, pH 7.9). Each treatment was added 50 μl of particle suspension, 50 μl of CaCl2 (2.5 M) and 20 μl of spermidine (0.1 M; free base). The suspensions were briefly sonicated, centrifuged for 10 min at 12,000 g, and the sediment was washed with 400 μl absolute ethanol. The particles were centrifuged under the above conditions and resuspended in 60 μl absolute ethanol. 10 μl of this suspension of particles coated with the four different DNA were used in each bombardment event.

**Table 2.** Different DNA concentrations used in PIG bombardment.

| No. Treatment/<br>Shooting Pressure | No. of larvae used | TnGV DNA concentration | DNA concentration pTnGV101 repair vector | DNA concentration pU6-BbsI-RNA | DNA concentration pU6-BbsI-RNA |
| --- | --- | --- | --- | --- | --- |
| Control | 20 | 0 | 0 | 0 | 0 |
| 1, 175psi | 20 | 1 µg | 1 µg | 1 µg | 1 µg |
| 2, 175 psi | 20 | 2 µg | 3 µg | 3 µg | 3 µg |
| 3, 175 psi | 20 | 2 µg | 5 µg | 5 µg | 5 µg |
| 4, 160psi | 20 | 1 µg | 1 µg | 1 µg | 1 µg |
| 5, 160 psi | 20 | 2 µg | 3 µg | 3 µg | 3 µg |
| 6, 160 psi | 20 | 2 µg | 5 µg | 5 µg | 5 µg |

### Bombardment conditions for the PIG system

Third instar larvae of *T. ni* were used. The insects were placed in 35 mm Petri dishes at 40 cm from the filtration unit. They were subsequently shot by HE with a shooting pressure of 160 and 175 psi, under vacuum chamber conditions of 15.75 inHg, respectively. Individuals who survived the shoot were placed on artificial diet. The expression of Cas9 (vector pHS70/Cas9) was induced after the bombardment with a thermal shock of 37 °C for one hour (Gratz et al., 2013). After seven days, the bombardment larvae were analyzed as described in the analysis of the recombinant TnGV EGFP by Dot Blot section.

### Recombinant baculovirus TnGV by PCR analysis

The extraction of recombinant TnGV DNA was performed as described above. To detect the *egfp* reporter gene, oligonucleotides from homologous regions D-ORF5 5′CCGctgagCAAGGTAGCCTCGTCTATTGGAC 3′ and R-ORF7 5′CCGctcgagTTCTTCATCTTCAATATTATGTC3′ were used using the PCR Reagent System Kit (Invitrogen, Life Technologies Inc.). The reaction mixture was prepared with l00 ng of recombinant TnGV DNA, 200 M dNTP’s, MgCl2 3 mm, 100 ng of each primer, 2.5 U Taq DNA polymerase (Invitrogen) and reaction buffer up to a total volume of 50 μl. The amplification conditions were: 95 °C for 1 min, followed by 30 cycles of 95 °C for 30 s, 42 °C for 1 min and 72 °C for 2 min, ending amplification at 72 °C for 7 min. The amplified fragment of 743 bp was visualized by electrophoresis in 1% agarose gel.

### Analysis of the Recombinant TnGV EGFP by Dot Blot

Seven days after infection of *T. ni* larvae by bombardment, treated individuals were collected and analyzed. Each larva was suspended in 10 μL PBS, frozen in N2 liquid and homogenized. The suspension was kept in ice. The homogenates were then transferred to a Nylon Biodyne TM membrane (Gaithersburg, EE. USA). Overexpressed EGFP bacterial extract was used as positive control and uninfected *T. ni* larvae as negative control. The membrane with the homogenates and controls was subjected to vacuum for 1 hours to ensure protein binding to the membrane. The membrane was blocked with 5% skimmed milk prepared in Tris-EDTA buffer (Tris-HCl 10 mm and EDTA 5 mm, pH 7) in slow agitation for 1 hours. It was then incubated with a polyclonal anti-GFP antibody (1: 2500) overnight at 4 ° C. The membrane was washed five times in PBS buffer supplemented with 0.1% Tween 20. The membrane was re-incubated overnight with the secondary antibody, Anti-IgG (rabbit conjugated to HRP (horseradish peroxidase) (Enzo Life Sciences, Inc., Farmingdale, USA). USA) (1:2500). It was washed five times in PBS buffer. HRP activity was detected with the ImmobilionTM Western Kit (Millipore, Damstadt, Germany).

## Results

### Construction of the repair vector

The regions *ORFlef2/orf603* and *orf1629* of the vector pAcUW31 were replaced by regions *orf5* and *orf7* from the TnGV baculovirus. *orf5* and *orf7* were previously amplified from the DNA of the TnGV wild strain. The resulting repair vector was called pTnGV101. The proper insertion of *orf5* and *orf7* was corroborated by DNA extraction, obtaining a corresponding band of 8000 bp (Figure 1), and by sequencing of recombinant plasmid.

**Figure 1.**
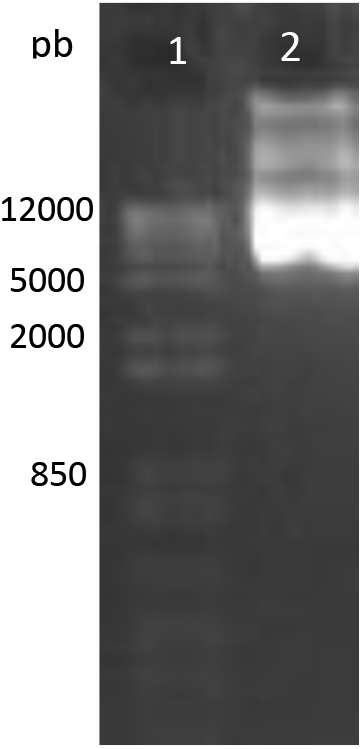
DNA of the pTnGV101 repair vector. Line 1: marker 1Kb Plus DNA Ladder (Invitrogen); line 2: 8000 bp band.

### Design and cloning of RNA guides in pU6-BBSI

The sgRNA (23pb) were designed and synthesized from the TnGV gene *p10*. Both sgRNA were successfully cloned at the BbsI site of plasmid pU6-BbsI, which expresses protein Cas9. The proper insertion of sgRNA at the pU6-BbsI cloning site was corroborated by PCR, obtaining a 700-850 bp amplicon (Figure 2), and by sequencing the modified plasmid pU6-BbsI-RNA.

**Figure 2.**
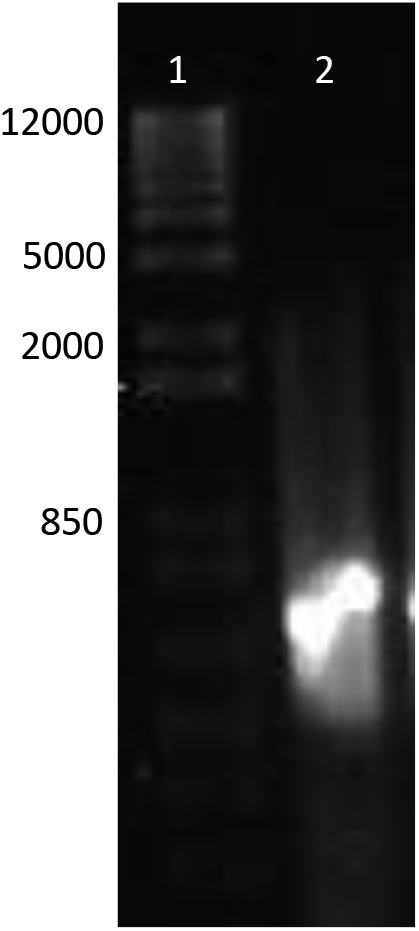
Amplification of the pU6-BbSI-RNA cloning site with gRNAs. Line 1: marker 1Kb Plus DNA Ladder (Invitrogen); Line 2: 850 bp.

### PIG co-transfection in *T. ni* larvae

Infection of *T. ni* larvae was achieved by bombardment with the PIG system using gold particles (60 mg/ml). With treatment 3 (see Tables 2 and 3), 175 psi of shooting pressure, 2 μg of TnGV DNA and 5 μg of DNA of pTnGV101, pU6-BBSI-RNA and pHS70/Cas9, respectively, 40% of the *T. ni* larvae bombardment were infected. Treatment 3 was followed by treatments 1, 2 (175 psi shooting pressure), 4, 6 and 5 (160 psi shooting pressure); with infection rates of 35, 30, 10, 10 and 5%, respectively (Table 3). Seven days after the bombardment of *T. ni* larvae, surviving individuals showed the characteristic infection caused by TnGV white body, slow movements (Figure 3A), while the control larvae were kept with green body and fully active (Figure 3B).

**Table 3.** *T. ni* larvae bombardment results using the PIG system.

| **No. Treatment/ Shooting Pressure** | **No. of larvae used** | **Dead larvae** | **Infected larvae** | **Infection rate** |
| --- | --- | --- | --- | --- |
| Control | 20 | 0 | 0 | 0% |
| 1, 175psi | 20 | 2 | 7 | 30% |
| 2, 175 psi | 20 | 3 | 6 | 30% |
| 3, 175 psi | 20 | 2 | 8 | 40% |
| 4, 160psi | 20 | 1 | 2 | 10% |
| 5, 160 psi | 20 | 2 | 1 | 5% |
| 6, 160 psi | 20 | 2 | 2 | 10% |

**Figure 3.**
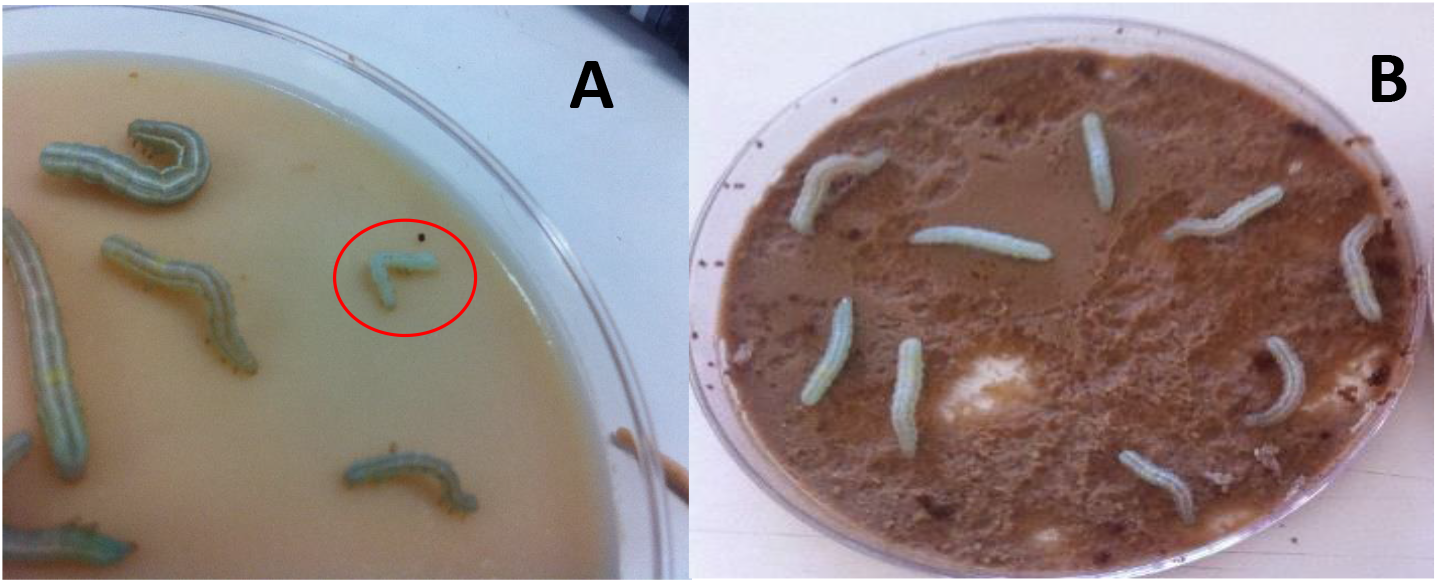
Co-transfection of *T. ni* larvae with recombinant TnGV. A. Larva of *T. ni* infected with recombinant TnGV (red circle). B. *T. ni* larvae or un-infected (negative control).

### Detection of EGFP in co- transfected larvae

PIG-infected *T. ni* larvae were processed for the detection of EGFP by Dot Blot estimating protein expression in 38% of individuals infected with the first three treatments (175 psi); and 10% of infected individuals in treatments 4, 5 and 6 (160 psi).

### Detection of the *egfp* gene by PCR

DNA was extracted from each infected larva with EGFP positive expression. A sample was subsequently selected for each pressure condition (2 samples) and amplification was performed for region *orf5* and *orf7*, with previously reported oligonucleotides. Two amplicons were obtained from 700 to 800 bp (Figure 5). The amplicons obtained were sequenced to corroborate the genetic modification in TnGV.

**Figure 5.**
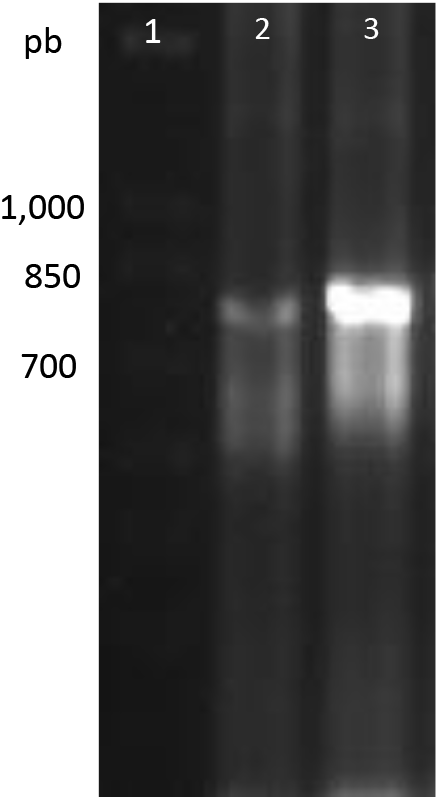
Amplification of *egfp* integrated into the genome of TnGV. Line 1: marker 1Kb Plus Ladder (Invitrogen); line 2: *egfp*, 160 psi; line 3: egfp,175 psi.

## Discussion

Baculovirus expression systems have had a profound impact on the production of recombinant proteins in insect cell lines (Hajós et al., 1998; Kost & Condreay, 1999; Luckow et al. 1993). However, cell lines that are permissible for baculovirus infections are not always available, limiting the collection of recombinant strains. This research demonstrated the potential to use the CRISPR/Cas9 system for genome editing of baculoviruses using the PIG bombardment method. This is very relevant as it allows the expression of heterologous genes in insect larvae.

The CRISPR/Cas9 system has recently been used for genetic modification in some large genome viruses (Ebina et al., 2013; King & Munger, 2019). A key part of the CRISPR/Cas9 system is the design of sgRNA (Suenaga et al., 2014). For this reason, there are different platforms for the *in silico* design of sgRNA. For this purpose, the CHOP-CHOP platform was used in this work. Testing sgRNA *in vivo* maintained the efficiency predicted by the program. In addition to sgRNA, there are other strategies such as dual sgRNA or multiplex that have been shown in previous studies to increase efficiency in genome editing (Chen et al., 2014).

The bombardment technique was also used to modify the TnGV genome. It was designed for plant transformation but has also been used to modify different types of organisms including insects (Sanford et al., 1995, Obregón-Barboza et al., 2007). In this work, the PIG method was used, with which a transformation efficiency of 40% and 38% was obtained, for two different pressure conditions in the modification of the genome of the baculovirus TnGV. Data similar have previously been reported by Obregón-Barboza et al., (2007). Their experiments consisted in testing different firing pressure conditions and type of microprojectiles (gold and tungsten) but, unlike our work, they used the PDS-1000/HE system, having favorable data for use in baculovirus.

In addition to the above, EGFP expression was very useful for detecting *T. ni* co-transfected larvae. Using the PCR and Dot Blot technique, the infected organisms were individually analyzed and the insertion of this reporter gene into the TnGV virus genome was verified. In other investigations *gfp* has been used as a reporter gene for the easy detection of genetic transformation of different types of organisms including insects as another of its advantages, is that it can be cloned into transfer vectors (Horn et al., 2002).

The use of two systems, the CRISPR/Cas9 and the bombardment PIG microprojectiles opens a wide variety of alternatives to recombinant viruses, not only to improve their virulence as biological pest control agents, but also as vectors of heterologous protein expression. Of course, this will require the development of standard protocols that should include the design of specific homologous repair vectors, the optimization of bombing conditions, and the greasy design. Theoretically, this technique has the potential to generate recombinant viruses using any baculovirus genus.

